# Transcriptomic Profiling Reveals Key Molecular Events and Pathways Driving Acquired Chemotherapy Resistance in Relapsed Acute Myeloid Leukemia

**DOI:** 10.64898/2025.12.24.696467

**Authors:** Min Zou, Jiaru Shi, Jing Hu, Chengming He, Min Zou

## Abstract

**Background:** Acute myeloid leukemia (AML) is a highly heterogeneous hematologic malignancy. Although induction therapy induces remission in many patients, relapse and acquired chemoresistance remain the major causes of treatment failure. Defining the molecular mechanisms underlying relapse is essential for improving therapeutic strategies.

**Methods:** Bone marrow samples from nine AML patients were analyzed, including five newly diagnosed cases and four relapsed cases after chemotherapy. Transcriptome sequencing and integrative bioinformatics analyses were performed, including differential expression analysis, GO/KEGG enrichment, GSEA, and protein–protein interaction (PPI) network analysis, to delineate relapse-associated molecular alterations.

**Results:** Principal component analysis demonstrated clear transcriptional segregation between primary and relapsed AML, indicating extensive molecular reprogramming during relapse. A total of 2,025 differentially expressed genes were identified, enriched in pathways related to epithelial–mesenchymal transition–like programs, leukemia stem cell maintenance, apoptosis evasion, and bone marrow microenvironment remodeling. Marked upregulation of FOXC1, HOXA11/HOXA11-AS, and AXL suggests key roles in sustaining stemness and promoting drug resistance. GO/KEGG analysis revealed coordinated activation of small GTPase, Rho/Ras signaling, ion transport, and epigenetic regulatory pathways, reflecting multilayered adaptive responses. GSEA indicated metabolic–epigenetic reprogramming in relapsed AML, while primary AML showed enrichment of energy metabolism and chromatin assembly pathways. PPI network analysis highlighted a central inflammation–metabolism axis involving TP53, IL6, CXCL8, and CCL2, associated with apoptosis resistance and metabolic adaptation.

**Conclusions:** Relapsed AML is characterized by transcriptional reprogramming, metabolic remodeling, and microenvironment-driven adaptive resistance. Targeting small GTPase signaling, AXL, or IL6-related inflammatory pathways, alone or combined with epigenetic modulators, may offer promising therapeutic strategies to overcome chemoresistance in AML.

## 1. Introduction

Acute myeloid leukemia (AML) is a group of clonal malignant neoplasms originating from hematopoietic stem/progenitor cells, characterized by abnormal proliferation of immature myeloid cells in the bone marrow and suppression of hematopoietic differentiation [1–3]. Although current standard induction chemotherapy (such as cytarabine combined with anthracyclines) can induce complete remission in most patients, the relapse rate remains as high as 50–70%, and the prognosis for relapsed patients is extremely poor [4–6]. Acquired chemotherapy resistance has become the primary obstacle to AML treatment failure, making in-depth analysis of its molecular mechanisms crucial for improving therapeutic strategies and enhancing patient survival rates [7,8].

Existing studies indicate that chemotherapy resistance in AML is not limited to upregulation of drug efflux pumps or abnormal drug metabolism but also involves the survival maintenance of leukemia stem cells (LSCs), the formation of epithelial-mesenchymal transition (EMT)-like phenotypes, and interactions with the bone marrow microenvironment [9–11]. For instance, LSC populations exhibit high self-renewal capacity and quiescence, conferring inherent resistance to cell cycle-specific chemotherapeutic agents [12]. Additionally, upregulation of receptor tyrosine kinases such as AXL has been demonstrated to promote resistance in FLT3-ITD+ AML by activating PI3K/AKT and MAPK pathways to enhance anti-apoptotic capabilities [13,14].

Meanwhile, in recent years, metabolic reprogramming and epigenetic remodeling have been recognized as important regulatory axes in resistance formation. Studies show that AML cells during relapse can adapt to chemotherapy-induced oxidative stress and DNA damage by enhancing oxidative phosphorylation (OXPHOS) activity, modulating NAD⁺/NADH ratios, and altering histone modification states [15–17]. Inflammatory factors in the bone marrow microenvironment (such as IL-6 and CXCL8) have also been confirmed to promote LSC survival and chemotherapy resistance through activation of STAT3/NF-κB signaling pathways [11,18].

Based on the above background, this study employs a longitudinal design to compare bone marrow samples from newly diagnosed and relapsed AML patients. Through transcriptome sequencing, differential expression analysis, GO/KEGG enrichment, GSEA, and protein-protein interaction (PPI) network analysis, we systematically investigate key molecular events and signaling pathways associated with AML chemotherapy relapse. Our research aims to reveal resistance mechanisms involving epigenetic remodeling, metabolic adaptation, and inflammation-microenvironment signaling interactions in relapsed AML, and to identify potential therapeutic targets. Through these analyses, we hope to provide new theoretical foundations and intervention strategies for preventing AML relapse and overcoming resistance.

## 2. Materials and Methods

### 2.1. Experimental Procedure

#### 2.1.1. Sample Quality Control

RNA integrity was assessed using the RNA Nano 6000 Assay Kit of the Bioanalyzer 2100 system (Agilent Technologies, CA, USA).

#### 2.1.2. Library preparation for Transcriptome sequencing

Total RNA was used as input material for the RNA sample preparations. Briefly, mRNA was purified from total RNA using poly-T oligo-attached magnetic beads. Fragmentation was carried out using divalent cations under elevated temperature in First Strand Synthesis Reaction Buffer(5X). First strand cDNA was synthesized using random hexamer primer and M-MuLV Reverse Transcriptase (RNase H-). Second strand cDNA synthesis was subsequently performed using DNA Polymerase I and RNase H. Remaining overhangs were converted into blunt ends via exonuclease/polymerase activities. After adenylation of 3’ ends of DNA fragments, Adaptor with hairpin loop structure were ligated to prepare for hybridization. In order to select cDNA fragments of preferentially 370∼420 bp in length, the library fragments were purified with AMPure XP system. Then PCR was performed with Phusion High-Fidelity DNA polymerase, Universal PCR primers and Index (X) Primer. At last, PCR products were purified (AMPure XP system) and library quality was assessed on the Agilent Bioanalyzer 2100 system.

#### 2.1.3. Clustering and sequencing (Novogene Experimental Department)

The clustering of the index-coded samples was performed on a cBot Cluster Generation System using TruSeq PE Cluster Kit v3-cBot-HS (Illumia) according to the manufacturer’s instructions. After cluster generation, the library preparations were sequenced on an Illumina Novaseq platform and 150 bp paired-end reads were generated.

### 2.2. Data Analysis

#### 2.2.1. Quality control

Raw data (raw reads) of fastq format were firstly processed through fastp software. In this step, clean data (clean reads) were obtained by removing reads containing adapter, reads containing ploy-N and low quality reads from raw data. At the same time, Q20, Q30 and GC content the clean data were calculated. All the downstream analyses were based on the clean data with high quality.

#### 2.2.2. Reads mapping to the reference genome

Reference genome and gene model annotation files were downloaded from genome website directly. Index of the reference genome was built using Hisat2 v2.0.5 and paired-end clean reads were aligned to the reference genome using Hisat2 v2.0.5. We selected Hisat2 as the mapping tool for that Hisat2 can generate a database of splice junctions based on the gene model annotation file and thus a better mapping result than other non-splice mapping tools.

#### 2.2.3. Novel transcripts prediction

The mapped reads of each sample were assembled by StringTie (v1.3.3b) (Mihaela Pertea.et al. 2015) in a reference-based approach. StringTie uses a novel network flow algorithm as well as an optional de novo assembly step to assemble and quantitate fulllength transcripts representing multiple splice variants for each gene locus.

#### 2.2.4. Quantification of gene expression level

FeatureCounts v1.5.0-p3 was used to count the reads numbers mapped to each gene. And then FPKM of each gene was calculated based on the length of the gene and reads count mapped to this gene. FPKM, expected number of Fragments Per Kilobase of transcript sequence per Millions base pairs sequenced, considers the effect of sequencing depth and gene length for the reads count at the same time, and is currently the most commonly used method for estimating gene expression levels.

#### 2.2.5. Differential expression analysis

(For DESeq2 with biological replicates) Differential expression analysis of two conditions/groups (two biological replicates per condition) was performed using the DESeq2 R package (1.20.0). DESeq2 provide statistical routines for determining differential expression in digital gene expression data using a model based on the negative binomial distribution. The resulting P-values were adjusted using the Benjamini and Hochberg’s approach for controlling the false discovery rate. Genes with an adjusted P-value <=0.05 found by DESeq2 were assigned as differentially expressed.

(For edgeR without biological replicates) Prior to differential gene expression analysis, for each sequenced library, the read counts were adjusted by edgeR program package through one scaling normalized factor. Differential expression analysis of two conditions was performed using the edgeR R package (3.22.5). The P values were adjusted using the Benjamini & Hochberg method. Corrected P-value of 0.05 and absolute foldchange of 2 were set as the threshold for significantly differential expression.

#### 2.2.6. GO and KEGG enrichment analysis of differentially expressed genes

Gene Ontology (GO) enrichment analysis of differentially expressed genes was implemented by the clusterProfiler R package, in which gene length bias was corrected. GO terms with corrected P value less than 0.05 were considered significantly enriched by differential expressed genes. KEGG is a database resource for understanding high-level functions and utilities of the biological system, such as the cell, the organism and the ecosystem, from molecular-level information, especially large-scale molecular datasets generated by genome sequencing and other high-through put experimental technologies (). We used clusterProfiler R package to test the statistical enrichment of differential expression genes in KEGG pathways.

#### 2.2.7. Gene Set Enrichment Analysis

Gene Set Enrichment Analysis (GSEA) is a computational approach to determine if a predefined Gene Set can show a significant consistent difference between two biological states. The genes were ranked according to the degree of differential expression in the two samples, and then the predefined Gene Set were tested to see if they were enriched at the top or bottom of the list. Gene set enrichment analysis can include subtle expression changes. We use the local version of the GSEA analysis tool, GO, KEGG data set were used for GSEA independently.

#### 2.2.8. PPI analysis of differentially expressed genes

PPI analysis of differentially expressed genes was based on the STRING database, which known and predicted Protein-Protein Interactions.

## 3. Results

### 3.1. Patient Grouping and Transcriptome Sequencing Data Quality Control

This study aims to explore the potential resistance mechanisms underlying chemotherapy relapse in acute myeloid leukemia (AML). To this end, we collected bone marrow samples from 9 patients and divided them into two independent groups for comparative analysis: 5 newly diagnosed AML (AML) patients and 4 relapsed AML (R_AML) patients post-chemotherapy. Among them, the basic clinical characteristics and treatment histories of 4 patients with poor-prognosis relapsed/refractory AML are as follows:

Patient 1 (R_AML_1) was first diagnosed on August 18, 2023, achieved complete remission after cytarabine combined with daunorubicin induction chemotherapy, and subsequently received consolidation therapy with daunorubicin + cytarabine, homoharringtonine, mitoxantrone + cytarabine + etoposide, and homoharringtonine regimens. The patient relapsed in February 2025, failed to achieve remission after cytarabine + doxorubicin + granulocyte colony-stimulating factor chemotherapy, and had an overall survival of 2 years post-diagnosis.

Patient 2 (R_AML_2) was first diagnosed on November 28, 2024, with initial treatment of venetoclax combined with azacitidine, achieving remission after 1 cycle, and completing 3 cycles before relapsing in June 2025. The patient is currently surviving with disease, with a survival time of 11 months post-diagnosis.

Patient 3 (R_AML_3) was first diagnosed on June 17, 2024, achieved remission after idarubicin combined with cytarabine induction chemotherapy, and subsequently received 5 cycles of idarubicin + cytarabine and cytarabine regimens. The patient relapsed in August 2025, failed to achieve remission after 2 cycles of cladribine + cytarabine + granulocyte colony-stimulating factor treatment, and has survived 16 months post-diagnosis.

Patient 4 (R_AML_4) was first diagnosed on December 27, 2022, achieved remission with the initial DA regimen, and achieved remission again after decitabine treatment but subsequently discontinued therapy. Relapsed on August 15, 2023, and sequentially received homoharringtonine + azacitidine + venetoclax, cytarabine, fludarabine + cytarabine + granulocyte colony-stimulating factor, and various combination chemotherapy regimens, but the disease did not remit, ultimately dying in July 2024 with an overall survival of 19 months post-diagnosis.

In summary, all 4 patients achieved short-term remission after initial treatment but experienced relapse or resistance in subsequent therapies, exhibiting varying survival differences. These samples provide a clinical foundation for subsequent differential gene and functional enrichment analyses and are representative. This longitudinal grouping strategy (relapsed vs. newly diagnosed) enables direct identification of molecular changes associated with acquired chemotherapy resistance. All samples successfully underwent RNA extraction post-sampling.

Subsequently, we performed transcriptome sequencing on all 9 samples. The sequencing output was stable, with an average of approximately 6.0 GB of raw data per sample. Sequencing quality was assessed for all samples, and key quality control metrics met analysis requirements (Table 1). High-quality data ensured the reliability of downstream bioinformatics analyses such as differential expression analysis.

**Table 1.** Summary of data quality.

In AML, acquired resistance often involves selection pressure induced by chemotherapy (such as cytarabine combined with anthracyclines), leading to the expansion of residual leukemia stem cells (LSCs) or resistant clones. These clones may maintain survival and self-renewal through intrinsic signaling pathway activation or microenvironment interactions. The longitudinal design helps capture dynamic transcriptome changes from diagnosis to relapse, such as how LSC localization in the bone marrow microenvironment enhances resistance via cell adhesion molecules (e.g., integrins).

In conclusion, the patient cohort and transcriptome sequencing data in this study provide a reliable foundation for exploring AML chemotherapy resistance mechanisms, revealing potential molecular dynamic changes through longitudinal comparison, and laying high-quality data support for subsequent analyses.

### 3.2. Differential Expression Gene Analysis Reveals Significant Transcriptomic Remodeling Associated with Relapse/Resistance

To assess the overall transcriptome differences between the two groups, we first performed principal component analysis (PCA). Samples from the newly diagnosed group (AML) and relapsed group (R_AML) showed a clear separation trend in two-dimensional space, with principal component 1 (PC1) and principal component 2 (PC2) explaining 23.82% and 18.75% of the total variance, respectively (Figure 1.A). This indicates significant transcriptomic heterogeneity in our sample grouping, particularly with the R_AML group distinctly separated from the AML group on PC1, suggesting substantial molecular expression differences between diagnostic and relapsed states.

**Figure 1.**
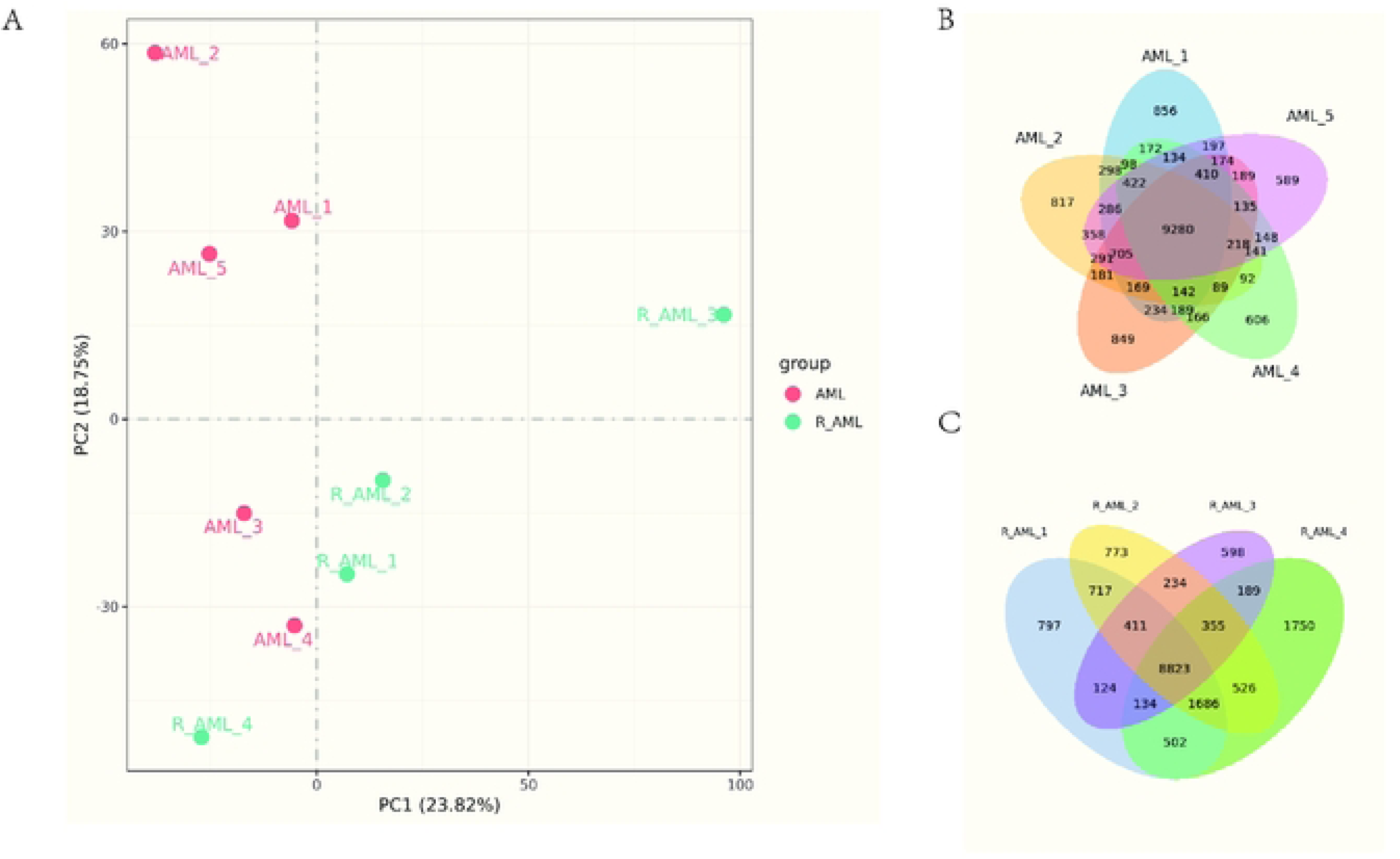
Quantitative analysis of samples. A. Principal component analysis (PCA) was performed to evaluate intergroup differences and within-group sample reproducibility. PCA was conducted using linear algebraic methods based on gene expression values (FPKM) across all samples. B, C. Venn diagrams showing the co-expressed genes in the AML and R_AML groups, respectively. The unique regions indicate genes expressed exclusively in each sample, while the overlapping areas represent genes commonly expressed across two or more samples.

To directly investigate molecular changes associated with chemotherapy resistance, we compared the relapsed group (R_AML) with the newly diagnosed group (AML). Using thresholds of |log2FC| ≥ 1 and adjusted p-value (padj) ≤ 0.05, we identified 2025 differentially expressed genes (DEGs), including 772 significantly upregulated genes and 1253 significantly downregulated genes(Figure 2.A). This extensive expression change strongly indicates that acquired chemotherapy resistance is accompanied by profound transcriptomic reconfiguration.

**Figure 2.**
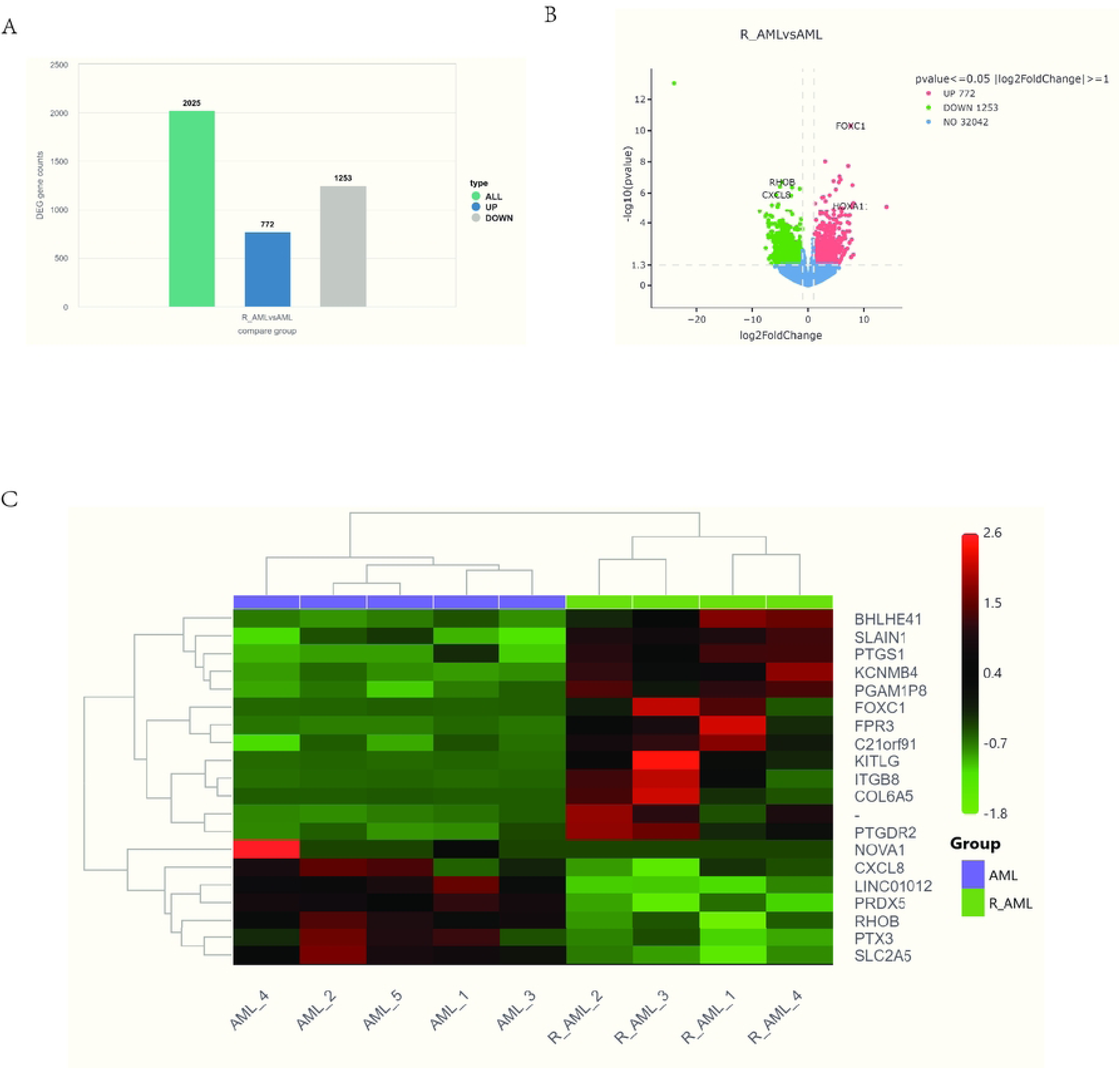
Differential gene expression analysis. A. Bar plot showing the number of differentially expressed genes (DEGs) between comparison groups, identified by DESeq2 with thresholds of p value ≤ 0.05 and |log₂FoldChange| ≥ 0.0. B. Volcano plot of DEGs. The x-axis represents log₂FoldChange values, and the y-axis represents −log₁₀(p value). Blue dashed lines indicate the threshold lines used for DEG selection. C. Hierarchical clustering heatmap of DEGs. The x-axis denotes sample names, and the y-axis shows normalized FPKM values of the DEGs.

Through comparative transcriptome analysis of relapsed AML (R_AML) and newly diagnosed AML (AML) samples, we identified a large number of significantly differentially expressed genes. These DEGs are primarily enriched in key pathways such as epithelial-mesenchymal transition (EMT)-like processes, leukemia stem cell (LSC) maintenance, apoptosis escape, inflammatory signaling, and bone marrow microenvironment interactions, collectively forming a complex resistance network.

In promoting EMT-like transition and LSC maintenance, several key genes exhibited significant upregulation(Figure 2.B,C). FOXC1 (log2FoldChange = 7.55, p = 4.92e-11), as a forkhead box transcription factor, enhances the malignant phenotype of leukemia cells by derepressing genes related to cell motility, invasion, and pro-survival pathways while suppressing apoptotic signals [19]. Its upregulation is closely associated with high HOX gene activity, collectively promoting LSC characteristic maintenance and the formation of protective bone marrow microenvironments [20]. Studies show that high FOXC1 expression is significantly correlated with reduced overall survival and event-free survival in patients, inducing treatment failure by upregulating matrix metalloproteinases (e.g., MMP7) and further enhancing cell metastatic potential and drug escape capabilities [21]. Synergizing with FOXC1 is the coordinated upregulation of HOXA11 (log2FoldChange = 7.76, p = 7.32e-06) and its antisense lncRNA HOXA11-AS (log2FoldChange = 7.23, p = 3.48e-05). This molecular axis expands EMT-like effects by suppressing p53-mediated apoptosis and promoting LSC self-renewal. Experiments confirm that HOXA11 knockdown enhances apoptosis via CD123/CD47 regulation and modulates NF-κB-related genes, restoring sensitivity to cytarabine [22]. HOXA11-AS may amplify this effect by stabilizing HOXA11 or silencing its inhibitors, similar to its role in promoting matrix metalloproteinase expression in other cancers. The upregulation of receptor tyrosine kinase AXL (log2FoldChange = 3.50, p = 3.77e-05) further reinforces this network. AXL enhances EMT-like transition, anti-apoptotic signals (e.g., BCL-2 upregulation), and LSC maintenance through activation of PI3K/AKT and MAPK pathways [23]. Notably, chemotherapy can induce AXL expression, mediating immune suppression and FLT3 signaling bypass, which is closely related to poor prognosis in resistant AML patients [24]. The high upregulation of long non-coding RNA XIST (log2FoldChange = 14.28, p = 7.80e-06) functions as a miRNA sponge (e.g., miR-142-5p, miR-124), elevating oncogene expression such as MYC and HMGB1, thereby suppressing apoptosis and enhancing stem cell characteristics [25].

In terms of microenvironment interactions and inflammatory signaling, we observed expression changes in a series of key genes. Downregulation of RHOB (log2FoldChange = −4.73, p = 1.99e-07) may promote cell adhesion-mediated drug resistance (CAM-DR) through integrin signaling and cytoskeletal remodeling. Downregulation of PTX3 (log2FoldChange = −5.10, p = 3.84e-07), encoding pentraxin-3, promotes M2 macrophage polarization via Akt/NF-κB signaling, forming an immunosuppressive microenvironment supportive of leukemia cell survival. Meanwhile, downregulation of CXCL8 (encoding IL-8, log2FoldChange = - 5.78, p = 1.30e-06) alters paracrine signaling from stromal cells, thereby upregulating BCL-2 and ERK1/2 pathways, supporting AML cell proliferation and inducing resistance to etoposide and other drugs [26]. Downregulation of PLAUR (encoding uPAR, log2FoldChange = −3.18, p = 1.63e-05) enhances CAM-DR via PI3K/AKT signaling, promoting cell migration and survival in the microenvironment [27].

Additionally, we identified expression changes in some novel transcripts (e.g., novel.1929), whose functions remain unclear but may participate in the aforementioned resistance network by influencing RNA processing or epigenetic regulation near known resistance loci.

In summary, differentially expressed genes in relapsed AML drive chemotherapy resistance and disease relapse through synergistic actions in EMT/LSC maintenance and microenvironment/inflammatory signaling. These genes provide a new theoretical basis for developing targeted therapeutic strategies (e.g., AXL inhibitors) [28].

### 3.3. GO Functional Enrichment Analysis Reveals Multi-Mechanism Synergy in Acquired Chemotherapy Resistance in Relapsed AML

Gene Ontology (GO) enrichment analysis revealed highly related signaling pathways significantly enriched in chemotherapy resistance, including small GTPase-mediated signal transduction (GeneRatio: 13/338, p-value: 0.007), regulation of Rho protein signal transduction (GeneRatio: 8/338, p-value: 0.017), Ras protein signal transduction (GeneRatio: 8/338, p-value: 0.038), and regulation of small GTPase-mediated signal transduction (GeneRatio: 9/338, p-value: 0.028)(Figure 3.A-C). These results suggest that upregulation of certain guanine nucleotide exchange factors (GEFs, e.g., FGD6) [29] may enhance GTP-binding activity of small GTPases (e.g., Ras, Rho, and Rap), maintaining sustained activation of downstream MAPK/ERK and PI3K/AKT pathways, thereby associating with epithelial-mesenchymal transition (EMT), cytoskeletal remodeling, tumor stem cell characteristic maintenance, and anti-apoptotic signaling, implying enhanced tolerance to drugs such as cisplatin or paclitaxel; for example, in small GTPase pathways, RAB25 has been shown in solid tumors (e.g., pancreatic and ovarian cancer) to participate in chemotherapy resistance by regulating vesicle transport and cell polarity changes [30], suggesting similar mechanisms may partially exist in hematologic malignancies, while Rho signaling mechanisms involve phosphorylation cascades of the RhoA-ROCK axis, regulating actin polymerization and stress fiber formation, potentially related to EMT and tumor microenvironment remodeling, and have been observed in various cancers associated with chemotherapy and radiotherapy resistance [31]; meanwhile, Ras pathway mechanisms focus on GTP hydrolysis defects in KRAS mutants, often leading to persistent activation of downstream RAF/MEK/ERK signaling in solid tumors, inducing adaptive resistance such as bypassing drug inhibition via PI3K pathway activation, with rapid resistance to KRAS G12C inhibitors in clinical trials often stemming from these GEF-mediated feedback loops [32], and similar adaptive feedback may also exist in AML’s Ras pathways; these GTPase pathways further interact with metal ion transport (GeneRatio: 8/338, p-value: 0.041) and transmembrane transporter activity/cation transporter activity (GeneRatio: 17/536, p-value: 0.001), where upregulated transporters (e.g., KCNMB4) may enhance ion efflux (e.g., iron, sodium, calcium, and copper), suggesting cells indirectly influence drug accumulation and redox balance by regulating metal ion homeostasis, thereby participating in chemotherapy resistance, while downregulated influx proteins (e.g., SLC31A2) further imply restrictions on drug uptake, forming synergistic resistance mechanisms, including abnormal copper transport potentially activating redox imbalance and ROS production, with metal ion transport abnormalities associated with tumor progression and resistance in gastric cancer [33]; simultaneously, these signaling and transport pathways may be regulated by epigenetic processes such as chromatin assembly (GeneRatio: 5/338, p-value: 0.016) and nucleosome assembly (GeneRatio: 5/338, p-value: 0.016), where downregulated linker histones (e.g., HIST1H1E) may lead to chromatin relaxation, enhancing transcriptional accessibility of certain genes, thereby providing an epigenetic basis for activation of GTPase and transport pathways, with mechanisms involving defects in replication-coupled nucleosome assembly, such as inhibition of the CAF-1 complex enhancing chromatin accessibility, allowing transcription factor binding to promoter regions to activate resistance-related genes, manifesting as epigenetic reprogramming-driven adaptive resistance in BRAF inhibitor-resistant melanoma [34]. Overall, these results suggest that small GTPase-related signaling pathways may serve as upstream regulatory nodes, forming multi-layered resistance regulatory networks through interactions with transport systems and epigenetic programs. Recent clinical trials show that combining Rho/Ras inhibitors (e.g., ROCK blockers) with epigenetic drugs (e.g., HDAC inhibitors) and transport modulators (e.g., nanoparticle-targeted SLC proteins) can synergistically reverse multidrug resistance, enhance chemotherapy sensitivity, and improve patient prognosis [35].

**Figure 3.**
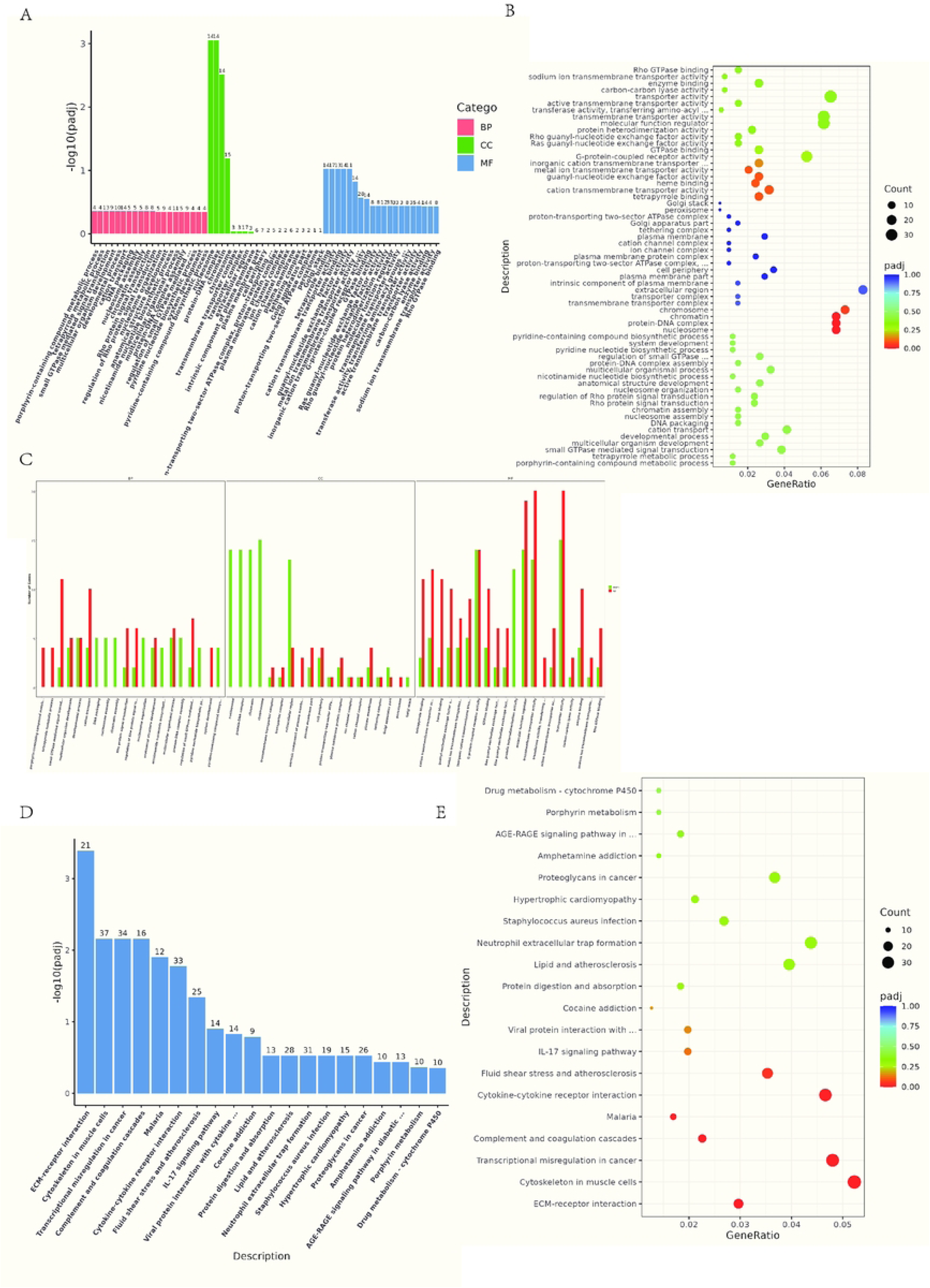
Functional enrichment analysis of differentially expressed genes. A. GO enrichment bar plot. The x-axis represents GO terms, and the y-axis shows enrichment significance, expressed as −log₁₀(padj). Different colors represent different functional categories. B. GO enrichment bubble plot. The x-axis represents the ratio of DEGs annotated to each GO term relative to the total number of DEGs, and the y-axis indicates the GO terms. Bubble size corresponds to the number of annotated genes, and color gradients from red to purple represent enrichment significance. C. Bar chart of up- and down-regulated GO terms. The x-axis shows the ratio of DEGs annotated to each GO term relative to the total number of DEGs, and the y-axis lists the corresponding GO terms. D. KEGG enrichment bar plot. The x-axis represents KEGG pathways, and the y-axis denotes enrichment significance (padj < 0.05 was considered significant). E. KEGG enrichment bubble plot. The x-axis represents the ratio of DEGs annotated to each KEGG pathway, and the y-axis lists the KEGG pathways. Bubble size indicates the number of annotated genes, and color gradients reflect the enrichment significance level.

### 3.4. KEGG Pathway Enrichment Analysis Identifies ECM Remodeling, Cytoskeletal Dynamics, and Immune Modulation as Key Drivers of Chemoresistance in Relapsed AML

To systematically elucidate the resistance mechanisms in relapsed AML, we performed KEGG pathway enrichment analysis on differentially expressed genes. The analysis revealed significant changes in pathways related to extracellular matrix (ECM) remodeling, cytoskeletal dynamics, transcriptional dysregulation, immune modulation, and drug metabolism (adjusted P-value < 0.05)(Figure 3.D,E). These pathways provide insights into the potential mechanisms of AML chemotherapy resistance, where upregulated genes often promote cell adhesion, migration, and anti-apoptotic signaling, while downregulated genes may weaken treatment responses. Top-ranked pathways such as ECM-receptor interaction and cytoskeleton regulation-related pathways (Cytoskeleton in muscle cells, core involving actin polymerization and stress fiber formation) highlight how leukemia cell interactions with their microenvironment and intracellular structural adaptations confer survival advantages under chemotherapy stress. The following discusses these findings based on current research progress, emphasizing their roles in disease progression and treatment failure.

The ECM-receptor interaction pathway, as the most significantly enriched pathway, includes upregulated genes such as integrins like ITGB5 and ITGA2B, and laminins like LAMC1, with downregulated genes including collagens like COL9A3 and thrombospondins like THBS1. This indicates bone marrow microenvironment remodeling favoring leukemia cell survival. In AML, ECM components interact with receptors on leukemia stem cells (LSCs) such as CD44, enhancing resistance by promoting quiescence and reducing drug-induced apoptosis. This is consistent with reports of ECM–integrin/CD44 signaling maintaining LSC quiescence [36,37,38]. Additionally, ECM adhesion activates mechanotransduction pathways, leading to metabolic reprogramming and upregulation of self-renewal genes, thereby enriching resistant populations and correlating with poor AML prognosis.

Closely linked is the cytoskeleton regulation-related pathway, whose core, although annotated in muscle cells, involves general actin polymerization and stress fiber formation, aligning with cell adhesion-mediated drug resistance (CAM-DR) mechanisms in AML. Upregulated genes include ACTA2 and integrins, with downregulated genes like MYH11 and troponins. This pathway reflects how AML cytoskeletal remodeling affects drug efflux, cell migration, and bone marrow stromal adhesion. For example, related RAC1 overactivation is associated with FLT3 inhibitor resistance, evading chemotherapy-induced cytoskeletal disruption through phosphorylation of resistance proteins, thereby reinforcing the protective role of the aforementioned ECM pathway [39–41].

Similarly, the transcriptional dysregulation in cancer pathway involves upregulated oncogenes such as MYC and HOXA9, with downregulated tumor suppressors like BCL6 and PAX5. In AML, chromatin rearrangements and transcription factor amplification lead to gene expression dysregulation, promoting stemness and blocking differentiation, which is key to resistance. This pathway aligns with HOXA9/MYC-driven stemness maintenance and may involve lncRNA such as GAS6-AS2-mediated epigenetic transcriptional activation, further sustaining anti-apoptotic networks and synergizing with the aforementioned structural adaptation mechanisms to amplify resistance effects [42–44].

The complement and coagulation cascades pathway shows upregulated complement components like C1QC, with downregulated factors like PLG, indicating immune escape and pro-thrombotic states in resistant AML. Although direct roles of complement cascades in AML require validation, C1q upregulation is associated with immune escape. This pathway regulates LSC transport and survival, potentially promoting resistance by altering the bone marrow microenvironment, echoing enhanced cell adhesion by coagulation elements and expanding the aforementioned microenvironment-dependent mechanisms [45].

The cytokine-cytokine receptor interaction pathway displays upregulated chemokines like CCL2 and receptors like CCR6, with downregulated interleukins like IL8. Cytokine networks in AML maintain inflammation and stem cell microenvironments, driving tumorigenesis and resistance. For example, IL-1 and TNF-α mediate extrinsic signals, synergizing with intrinsic factors like CEBPB to block differentiation and promote survival. This chronic inflammatory signaling activation as an extrinsic resistance driver overlaps with the IL-17 signaling pathway, where IL-17A promotes cell proliferation via PI3K/Akt and JAK/STAT3 activation, and IL-17B-IL-17RB upregulates BCL-2 to enhance survival, thereby interweaving with transcriptional dysregulation pathways to reinforce immune regulatory imbalances [46,47].

The neutrophil extracellular trap (NET) formation pathway enrichment primarily reflects differential expression of histone-related genes. Although NETosis is related to tumor immune modulation, its role in AML requires further study and may only represent indirect signals of immune response changes. NETs formed by neutrophils can bind anthracyclines to reduce efficacy, while impaired NETosis alters the immune landscape to promote poor prognosis, synergizing with the aforementioned immune pathways to exacerbate resistance [48,49].

The proteoglycans in cancer pathway involves upregulated syndecans like SDC1 and integrins, promoting tumor migration and growth in AML. These macromolecules support resistance by regulating microenvironment signaling cascades, linking with the ECM pathway [50].

Finally, the drug metabolism - cytochrome P450 pathway shows upregulated CYPs like CYP3A7, with downregulated GSTs like GSTM3, directly impacting AML resistance. The CYP family regulates drug metabolism in the microenvironment, such as CYP3A4 inactivating cytarabine and daunorubicin to protect cells, thereby integrating with the aforementioned metabolic reprogramming mechanisms to form a comprehensive resistance network [51].

Other pathways such as malaria, fluid shear stress and atherosclerosis, and viral protein interaction with cytokines may reflect overlapping immune and inflammatory responses, while addiction-related pathways imply neurotransmitter-like signaling in resistance. Overall, these findings indicate that AML chemotherapy resistance arises from multifaceted interactions of genetic dysregulation, microenvironment cues, and metabolic adaptations. Targeting these pathways, such as with ECM disruptors or cytokine inhibitors, may enhance therapeutic efficacy, supported by current studies.

### 3.5. GSEA Enrichment Analysis Reveals Resistance Phenotypes in Relapsed AML Acquired Through Protein Modification, Epigenetic Remodeling, and Cytoskeletal Suppression

To further dissect the system-level transcriptional reprogramming features of AML chemotherapy relapse, we performed gene set enrichment analysis (GSEA) [52,53]. We delved into the systemic differences in transcriptional regulation and metabolic characteristics between relapsed AML and newly diagnosed AML. The results show that relapsed AML exhibits significant transcriptional reprogramming features, while newly diagnosed AML displays unique metabolic dependencies, providing new perspectives for understanding AML disease progression and resistance mechanisms.

In relapsed AML, significant enrichment of pathways such as RNA biosynthetic processes and nucleic acid-templated transcription reflects profound transcriptional reprogramming during disease progression(Figure 4.A). These transcriptional changes are closely related to epigenetic regulation, where dynamic changes in DNA methylation and histone modifications may drive the establishment of adaptive gene expression programs by affecting key transcription factor activities [54–56]. Particularly noteworthy is the enrichment of insulin-like growth factor binding processes, suggesting an important role of the IGF signaling axis in relapse. Studies indicate that IGF signaling not only maintains leukemia stem cell survival via the PI3K/AKT pathway but may also enhance cellular transcriptional plasticity by regulating epigenetic modification states, thereby promoting resistance phenotype formation [57,58].

**Figure 4.**
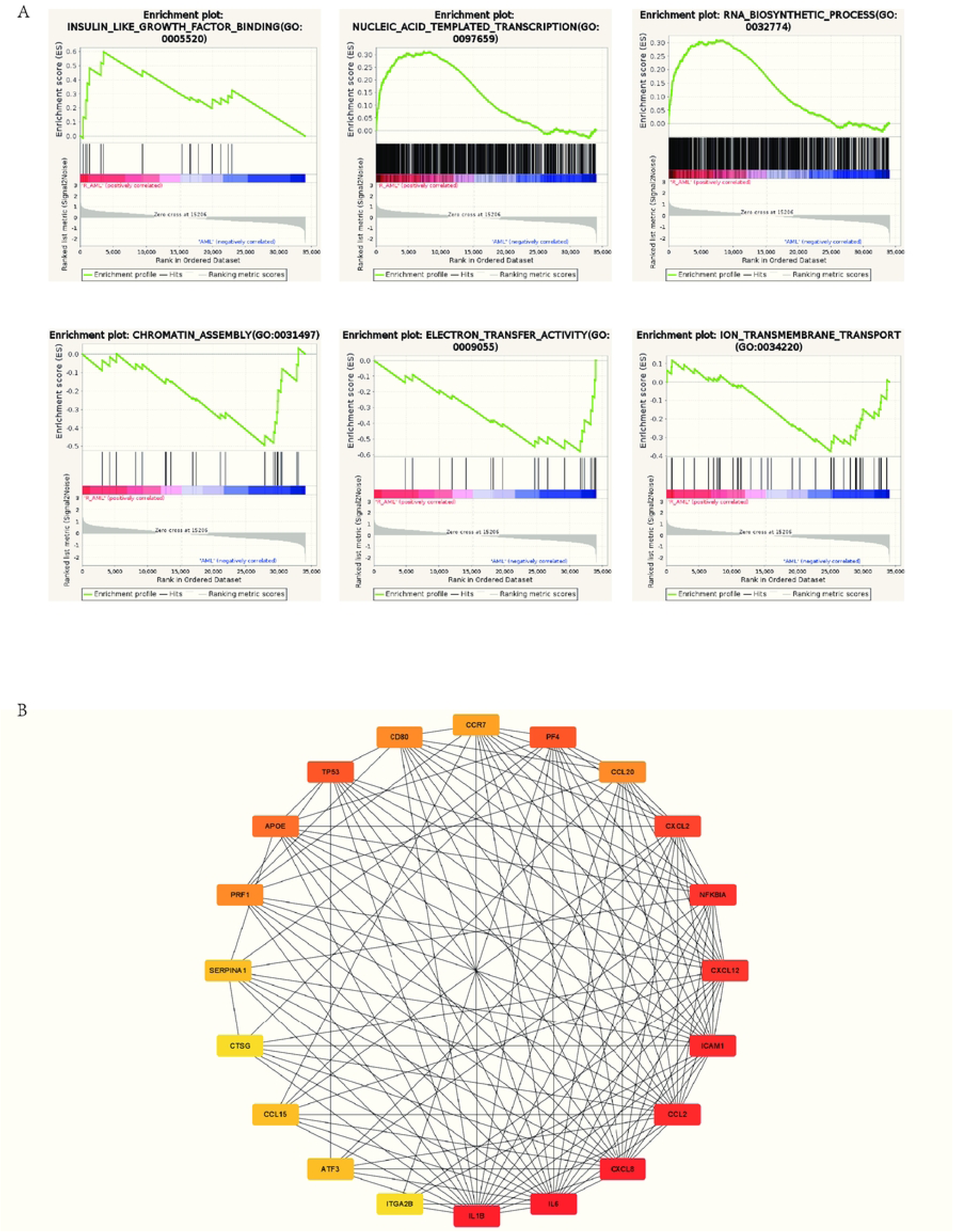
GSEA enrichment and protein–protein interaction (PPI) network analysis. A. Enrichment score (ES) line plot. The x-axis represents the ranked position of each gene within the gene set, and the y-axis indicates the running enrichment score (ES). The peak of the curve corresponds to the maximal enrichment score of the gene set. A positive normalized enrichment score (NES) indicates enrichment in the R_AML group, whereas a negative NES indicates enrichment in the newly diagnosed AML group. B. Protein–protein interaction (PPI) network. Each node represents a protein, and each edge denotes an interaction between connected proteins.

In stark contrast, newly diagnosed AML exhibits a characteristic pattern centered on energy metabolism and cellular structural assembly. Enrichment of pathways such as electron transfer activity, ion transmembrane transport, and chromatin assembly reflects the dependence of diagnostic leukemia cells on active metabolic states [59,60]. This metabolic feature may form the basis for cells’ initial responses to cytotoxic drugs. During disease relapse, leukemia cells undergo significant metabolic reprogramming, shifting from energy metabolism modes dominated by glycolysis and oxidative phosphorylation to alternative pathways such as fatty acid β-oxidation [61]. This metabolic adaptation not only reduces cellular energy consumption but may also indirectly influence epigenetic states and gene expression programs by altering intracellular metabolite compositions.

The interplay between transcriptional reprogramming and metabolic adaptation constitutes the core of AML relapse mechanisms. Epigenetic regulation serves as a key hub connecting these two levels, integrating signals from metabolic pathways through mechanisms such as histone modifications and DNA methylation, thereby regulating gene expression networks related to cell stemness maintenance and drug resistance [62,63]. For example, specific metabolites produced during metabolic reprogramming may alter chromatin states by affecting histone-modifying enzyme activities, ultimately promoting leukemia stem cell characteristic maintenance. This dynamic networked regulatory mechanism reveals the complexity of cellular state evolution from diagnosis to relapse in AML and provides a theoretical basis for developing novel therapeutic strategies targeting the epigenetic-metabolic axis.

### 3.6. PProtein Interaction Network Analysis Reveals the Inflammation-Metabolism Axis as a Core Regulatory Hub for Resistance in Relapsed AML

Protein-protein interaction (PPI) network analysis based on the STRING database revealed a network involving 56 unique proteins with 193 predicted interactions, with combined scores ranging from 0.403 to 0.999(Figure 4.B). Approximately 45% of edges have high confidence (combined score >0.900), suggesting strong functional or pathway associations among these proteins. The network exhibits a modular structure, with major clusters divisible into erythroid-related protein modules (possibly reflecting AML cells’ erythroid differentiation tendency or stress-induced expression features), inflammatory cytokine modules, and apoptosis regulation modules. This modular organization was identified using clustering algorithms in STRING’s network visualization tool.

Key hubs in the network include TP53 (p53), which shows predicted interactions with multiple partners such as ATF3 (combined score: 0.982), NFKBIA (0.915), and PMAIP1 (0.970), highlighting its potential role as a central tumor suppressor. Inflammatory mediators form another dense module, with CCL2 (MCP-1) tightly connected to CXCL8 (IL-8; 0.999), CXCL2 (0.929), and IL6 (0.985), reflecting coordinated chemokine signaling. Erythroid-related proteins, such as HBA2, HBG2, and AHSP, form tight clusters with scores exceeding 0.990, suggesting stress responses or AML cells’ erythroid differentiation features.

Quantitative analysis highlights enriched interaction types: cytokine pairs show elevated co-expression scores (e.g., CCL2-CXCL8: 0.994), while apoptosis-related edges are primarily dominated by database-annotated interactions (e.g., BCL2A1-PMAIP1: 0.907). The average node degree is 6.89, with TP53, CCL2, and CXCL8 having degrees >10, indicating their centrality in the predicted network.

Based on these predicted interaction features, the identified PPI network provides insights into potential chemotherapy resistance mechanisms in relapsed AML. The prominence of TP53 interactions aligns with TP53 mutations in AML cases, which are associated with anthracycline resistance by impairing DNA damage-induced apoptosis [64]. Existing studies indicate that pre-existing TP53 mutations may confer chemotherapy resistance and lead to selective expansion of mutant clones post-treatment, as these alterations disrupt activation of pro-apoptotic effectors like PMAIP1 (NOXA) [65]. For example, high-score predicted connections of TP53 with PMAIP1 (0.970) and BCL2A1 (0.699) suggest possible dysregulation of intrinsic apoptosis pathways. Research shows that reduced NOXA expression in TP53-mutant cells impairs mitochondrial outer membrane permeabilization, leading to hindered cytochrome c release and caspase activation, potentially evading doxorubicin-induced cell death [66]. Additionally, recent evidence suggests that TP53-mutant AML cells may rely on the mevalonate pathway for survival under cytarabine (AraC) stress, where inhibiting this pathway may restore sensitivity by exacerbating metabolic vulnerability [67].

The inflammatory module, including IL6, IL1B, and CXCL8, may promote resistance through the tumor microenvironment (TME). Elevated IL6 signaling (e.g., IL6-CXCL8: 0.991) is associated with STAT3 activation, potentially forming protective niches for leukemia blasts from cytarabine [68]. Some studies indicate that IL6 can induce MFN1-mediated mitochondrial fusion, thereby potentially enhancing oxidative phosphorylation (OXPHOS) and ATP production to maintain energy demands and confer apoptosis resistance during chemotherapy assaults [69]. Furthermore, IL6 may upregulate CD36-mediated fatty acid uptake in AML cells, further reinforcing lipid metabolism to counter drug-induced oxidative stress and promote survival [70]. Bone marrow mesenchymal stromal cells (MSCs) exacerbate this process by secreting IL6, with some studies suggesting that IL6 transfers functional mitochondria to AML cells via tunneling nanotubes, thereby enhancing OXPHOS and chemotherapy resistance through JAK2/STAT3 signaling [71]. Similarly, CXCL8 (IL-8), often co-expressed with IL6, may contribute to TME remodeling by recruiting immunosuppressive myeloid-derived suppressor cells (MDSCs) and promoting cancer cell plasticity, leading to acquired resistance to multiple chemotherapeutic agents [72].

CCL2-CXCR4 interactions (indirectly mediated via CXCL12; 0.841) can enhance bone marrow homing and quiescence, with mechanisms involving persistence of minimal residual disease post-chemotherapy [73]. CCL2, produced by MSCs or AML cells, may activate PI3K/Akt/mTOR signaling to inhibit apoptosis and mediate resistance. Existing models indicate that blocking CCL2 signaling can restore sensitivity to targeted therapies [74]. In the AML TME, CCL2 and CXCL8 may synergistically recruit MDSCs and regulatory T cells, creating an immunosuppressive barrier that weakens drug efficacy and promotes relapse [75].

The erythroid cluster proteins (e.g., AHSP-HBA2: 0.995) may reflect dysregulated oxidative stress response regulation rather than true erythroid differentiation. Because AML cells regulate erythroid genes under chemotherapy pressure to mitigate reactive oxygen species (ROS), thereby promoting survival [76]. AHSP binds alpha-hemoglobin (encoded by HBA2), inhibiting ROS production and protecting against oxidative damage, which can be hijacked in relapsed AML to counter chemotherapy-induced ROS overload [77]. Aberrant OXPHOS activation, potentially related to these proteins, has been shown to enhance treatment resistance in FLT3-mutant AML through metabolic adaptation and reduced apoptosis induction [78]. Although HBA1/HBA2 expression is often reduced in AML, leading to impaired growth inhibition of leukemia cells, the tight clusters in our network suggest compensatory upregulation in resistant clones to exploit ROS thresholds for survival. Overall, these predicted interactions suggest that relapsed AML may exploit inflammation, apoptosis, and metabolic dysregulation to evade treatment. This points to potential intervention directions, such as TP53 activators, IL6/JAK inhibitors, or ROS modulators, but further functional validation and clinical studies are needed.

## 4. Discussion

This study, through transcriptome sequencing analysis of bone marrow samples from newly diagnosed and relapsed/refractory acute myeloid leukemia (R_AML) patients, reveals molecular reprogramming processes closely associated with chemotherapy resistance and disease relapse. A total of 2025 differentially expressed genes (DEGs) were identified, with 772 upregulated and 1253 downregulated, indicating significant transcriptional level remodeling in AML under chemotherapy pressure. Principal component analysis (PCA) results show clear separation of diagnostic and relapsed group samples in overall expression profiles, suggesting systemic reconfiguration of global gene expression patterns in relapsed AML. These changes not only reflect accumulated mutation loads but also embody acquired resistance driven by transcriptional plasticity, metabolic reprogramming, and microenvironment adaptation under chemotherapy pressure.

At the molecular level, significant upregulation of genes such as FOXC1, HOXA11, AXL, and XIST in relapsed AML samples suggests that tumor cells acquire epithelial-mesenchymal transition (EMT)-like features, leukemia stem cell (LSC) maintenance capabilities, and anti-apoptotic phenotypes. FOXC1 and HOXA11 have been confirmed to maintain stemness and correlate with poor prognosis through activation of HOX-dependent transcriptional networks and promotion of extracellular matrix (ECM) remodeling [79,80]. High AXL expression aligns with its role in multiple tumors by mediating EMT and bypass signaling under chemotherapy stress via the PI3K/AKT pathway, thereby enhancing cell survival [81]. Upregulation of long non-coding RNA XIST may further promote resistance phenotype formation by regulating MYC and anti-apoptotic signaling pathways as a competing endogenous RNA (ceRNA) [82]. Conversely, downregulation of RHOB, PTX3, and CXCL8 suggests weakened inflammatory responses and immune cell recruitment functions, potentially leading to the formation of an immunosuppressive microenvironment in relapsed AML [83].

Functional enrichment analysis results further support the above inferences. GO analysis shows that small GTPase-mediated signal transduction, ion transmembrane transport, and chromatin assembly are significantly enriched biological processes in relapsed AML, suggesting cytoskeletal remodeling and nuclear reprogramming jointly promote cell migration and transcriptional plasticity [84]. KEGG analysis identifies activation of ECM-receptor interactions, focal adhesions, and cytokine signaling pathways, indicating that bone marrow microenvironment structural and signaling remodeling plays a key role in resistance formation [85]. GSEA results show that metabolic dependencies in relapsed AML shift from glycolysis in the diagnostic phase to RNA biosynthesis and epigenetic plasticity, suggesting relapsed cells possess stronger metabolic flexibility and transcriptional adaptability [86]. Protein-protein interaction (PPI) network analysis shows TP53, IL6, and CXCL8 as key network hubs, revealing the integrative roles of apoptosis imbalance, inflammatory responses, and metabolic adaptation in the resistance network.

These findings are consistent with previous research. Existing evidence indicates that FOXC1 and HOXA family genes promote resistance and relapse by maintaining leukemia stem cell self-renewal and quiescent states [79,87]; AXL signaling mediates immune escape and stemness maintenance post-chemotherapy [88]; while activation of the IL-6/JAK-STAT3 axis in the bone marrow microenvironment enhances residual leukemia cell survival [89]. Enrichment of small GTPase and cytoskeletal remodeling signals aligns with previous studies on the RhoA–ROCK pathway’s role in cell adhesion-mediated resistance (CAM-DR) in hematologic malignancies [90]. Enhancement of ECM-receptor and cytokine signaling pathways is consistent with protective mechanisms mediated by integrins, CXCL12–CXCR4, and IL-6 [91]. Enrichment of chromatin assembly and RNA processing pathways in GSEA also supports the core role of epigenetic reprogramming in acquired resistance [86], indicating that post-chemotherapy cells achieve transcriptional plasticity through histone modifications and chromatin remodeling to maintain survival.

Notably, some results differ from literature reports. For example, downregulation of CXCL8 in this study contrasts with its upregulation in AML progression in some studies [92], which may relate to negative feedback inhibition or clonal evolution after multiple chemotherapy rounds. Similarly, upregulation of erythroid-related genes (e.g., HBA2) in the PPI network may reflect stress-induced oxidative stress responses rather than true differentiation tendencies [93]. These differences may stem from variations in patient treatment histories, molecular subtypes, or sample compositions, suggesting significant heterogeneity in molecular mechanisms of AML relapse.

This study still has certain limitations. The small sample size (n=9) limits statistical power and generalizability of results; bulk RNA sequencing cannot resolve specific changes in different cell populations, and differences in blast proportions and stromal contamination in bone marrow may affect DEG identification. Although strict quality control was performed, batch effects were not completely eliminated. Future studies should expand sample sizes and use paired samples for validation in multi-center cohorts. Single-cell transcriptomics and multi-omics integration (e.g., proteomics, epigenomics) will help dissect resistant clone evolution and signaling network reconstruction. Validating the oncogenic roles and clinical potential of FOXC1, AXL, and IL6 signaling pathways through patient-derived xenograft (PDX) models or CRISPR-mediated gene editing is warranted.

In terms of clinical translation, the identified transcriptional features hold promise as biomarkers for relapse risk prediction and treatment stratification. FOXC1 and AXL may serve as potential therapeutic targets, with AXL inhibitors (e.g., bemcentinib) or strategies blocking ECM-integrin interactions potentially restoring drug sensitivity in resistant cells [94]. The inflammation-metabolism coupling axis revealed by the PPI network suggests disrupting resistant microenvironments through combined JAK/STAT pathway inhibitors and metabolic regulators (e.g., mitochondrial OXPHOS inhibitors) [95]. Further blocking CCL2/CCR2 or CXCL12/CXCR4 signaling may also weaken tumor-stroma protective interactions, thereby enhancing chemotherapy effects [96].

In summary, this study reveals a multi-dimensional resistance network in relapsed AML centered on EMT-like transcriptional reprogramming, leukemia stem cell maintenance, inflammation-metabolism coupling, and microenvironment remodeling, elucidating key molecular bases for chemotherapy failure and proposing multiple pathways with potential translational value. These results provide important theoretical and experimental foundations for further dissecting relapsed AML resistance mechanisms and developing combined targeted therapeutic strategies.

## Institutional Review Board Statement

The study was conducted in accordance with the Declaration of Helsinki, and approved by the Institutional Review Board (or Ethics Committee) of Ganzhou People’s Hospital(2022—ZD1368).

## Informed Consent Statement

Informed consent was obtained from all subjects involved in the study.

Written informed consent has been obtained from the patient(s) to publish this paper.

## Data Availability Statement

The original contributions presented in this study are included in the article/Supplementary Material. Further inquiries can be directed to the corresponding author.

